# Binocular Integration Manifests as a Transient Spiking Increase Followed by Selective Suppression in Primary Visual Cortex

**DOI:** 10.1101/318667

**Authors:** Michele A. Cox, Kacie Dougherty, Jacob A. Westerberg, Michelle S. Schall, Alexander Maier

## Abstract

Research throughout the past decades revealed that neurons in primate primary visual cortex (V1) rapidly integrate the two eyes’ separate signals into a combined binocular response. The exact mechanisms giving underlying this binocular integration remain elusive. One open question is whether binocular integration occurs at a single stage of sensory processing or in a sequence of computational steps. To address this question, we examined the temporal dynamics of binocular integration across V1’s laminar microcircuit of awake behaving monkeys. We find that V1 processes binocular stimuli in a dynamic sequence that comprises at least two distinct phases: A transient phase, lasting 50-150ms from stimulus onset, in which neuronal population responses are significantly enhanced for binocular stimulation compared to monocular stimulation, followed by a sustained phase characterized by widespread suppression in which feature-specific computations emerge. In the sustained phase, incongruent binocular stimulation resulted in response reduction relative to monocular stimulation across the V1 population. By contrast, sustained responses for binocular congruent stimulation were either reduced or enhanced relative to monocular responses depending on the neurons’ selectivity for one or both eyes (i.e., ocularity). These results suggest that binocular integration in V1 occurs in at least two sequential steps, with an initial additive combination of the two eyes’ signals followed by the establishment of interocular concordance and discordance.

**Significance Statement:** Our two eyes provide two separate streams of visual information that are merged in the primary visual cortex (V1). Previous work showed that stimulating both eyes rather than one eye may either increase or decrease activity in V1, depending on the nature of the stimuli. Here we show that V1 binocular responses change over time, with an early phase of general excitation and followed by stimulus-dependent response suppression. These results provide important new insights into the neural machinery that supports the combination of the two eye’s perspectives into a single coherent view.

## Introduction

The primate binocular visual field is the largest among mammals (Ross, 2000; Heesy, 2004; Heesy et al., 2011). Ocular convergence comes at the cost of total visual field size but improves visual perception in several ways: stereopsis allows for fine depth discrimination (Howard, 2012; Howard and Rogers, 2012), and binocular fusion can compensate for the retinal blind spot as well as some kinds of occlusion (Howard and Rogers, 2002). Furthermore, binocular viewing increases visual acuity and contrast sensitivity compared to viewing with one eye alone (Blake and Fox, 1973; Jones and Lee, 1981). These perceptual benefits of binocular vision require that inputs from the two eyes are rapidly integrated and combined into a singular coherent (“cyclopean”) view by the visual system. However, the two eyes’ views are almost never identical and often differ substantially (Wheatstone, 1838). Thus, the visual system must contain fast-acting mechanisms for detecting binocular concordance and resolving interocular differences to create a coherent view from output of the two eyes’ retinae (Cumming and DeAngelis, 2001; Tanabe and Cumming, 2014).

Anatomically, the retinogeniculate projections for each retina are largely segregated until converging onto single neurons within primary visual cortex (V1; Casagrande and Boyd, 1996). Geniculate axons from each eye terminate in horizontally alternating bands within V1’s granular layer (layer 4C in Old-World primates), giving rise to tangential regions of neurons that share eye preference, known as ocular dominance columns (Hubel and Freeman, 1977; Hendrickson et al., 1978). The functional consequence of this organization is that most V1 neurons respond more vigorously to stimulation in one eye compared to the other, though only a small minority of V1 neurons respond to one eye exclusively (Hubel and Wiesel, 1968; Kato et al., 1981; Wiesel, 1982; Malach et al., 1993; Smith et al., 1997). V1 neurons also exhibit tangentially organized selectivity for other stimulus features such as orientation (Hubel and Wiesel, 1968; Van Essen et al., 1984; Ringach et al., 1997). Given that its neurons encode both detailed stimulus features and eye-of-origin information, V1 is thought to play a critical role in detecting and resolving interocular differences (see Cumming and DeAngelis, 2001; Tong et al., 2006; Blake and Wilson, 2011; Read and Cumming, 2017 for review).

Since most V1 neurons receive inputs from both eyes, one might expect their synaptic activation to increase when both eyes are stimulated compared to stimulation of one eye alone. However, under some circumstances V1 neurons have been reported to *decrease* in spiking activity during binocular stimulation compared to monocular stimulation (Endo et al., 2000; Kumagami et al., 2000), an occurrence termed interocular suppression (Bishop et al., 1959; Vastala, 1960). A particularly well-studied variant of interocular suppression is dichoptic cross-orientation suppression (dCOS). In dCOS, a neuron’s response to an optimally oriented stimulus viewed by the neuron’s preferred eye is reduced by the presentation of an orthogonally oriented stimulus in the other eye (Sengpiel and Blakemore, 1994; Sengpiel et al., 1995a; 1995b; 1998; Sengpiel, 2005; Endo et al., 2000). The majority of dCOS studies averaged stimulus responses over long (>1s) exposure times. As a consequence, we know little about the temporal dynamics of interocular suppression and binocular integration.

The primary goal of this study was to examine the temporal dynamics of V1 binocular integration generally and dCOS specifically using matching (congruent) and non-matching (incongruent) binocular stimuli in awake primates. Our results suggest that there are at least two distinct phases of binocular computations in V1: An initial phase, roughly 150 ms in duration, characterized by response enhancement for binocular stimulation compared to stimulation of one eye alone. This initial binocular response enhancement is followed by a period of response reduction, matching previous reports of dCOS in cat area 17. Neurons exhibited interocular suppression regardless of their selectivity for one eye over the other (i.e. ocularity). However, neurons that responded equivalently to stimulation in each of the two eyes were uniquely excited by congruent binocular stimuli compared to both incongruent binocular stimulation and stimulation of one eye alone. Laminar analyses revealed that neurons in V1’s superficial layers, which are downstream of V1’s granular input layer, demonstrated the greatest response reduction for incongruent stimulation. These results suggest that the bulk of feature-specific computations underlying binocular integration occur beyond the granular input layer and are likely mediated by intracortical mechanisms.

## Materials and Methods

### Subjects

Two adult monkeys (*Macaca radiata*, one female) were used in this study. All procedures followed regulations by the Association for the Assessment and Accreditation of Laboratory Animal Care (AAALAC), were approved by Vanderbilt University’s Institutional Animal Care and Use Committee and observed National Institutes of Health guidelines.

### Surgical Procedures

In a series of surgeries, each monkey was implanted with a custom-designed MRI-compatible plastic head holder and a plastic recording chamber (Schmiedt et al., 2014). All surgeries were performed under sterile surgical conditions using isoflurane anesthesia (1.5–2.0%). Vital signs, including blood pressure, heart rate, SpO_2_, CO_2_, respiratory rate, and body temperature, were monitored continuously. During surgery, the head holder and a recording chamber were attached to the skull using transcranial ceramic screws (Thomas Recording) and self-curing denture acrylic (Lang Dental Manufacturing). A craniotomy was performed over the perifoveal visual field representation of primary visual cortex (both hemispheres in each monkey) concurrent with the position of the recoding chamber. Each monkey was given analgesics and antibiotics for post-surgical care.

### Visual Stimulation

Stimuli were presented on CRT monitors running at either 60 Hz (resolution 1280 × 1024 pixels; N = 7 sessions) or 85 Hz (resolution 1024 × 768; N = 23 sessions). CRT monitors were linearized by measuring the luminance of the monitor at 17 brightness increments, linearly spaced between the minimum and maximum for each gun (red, green, and blue), using either a photometer or spectroradiometer (PhotoResearch). We fit the measured luminance changes to a power function and then applied the inverse of the exponent of this power function for each R, G, and B values during stimulus generation. Visual stimuli were generated by MonkeyLogic (Asaad and Eskandar, 2008; Asaad et al., 2013) for MATLAB (R2012-2014, The MathWorks) running on a PC (Dell, Windows 7 or 10) with a NVIDIA graphics card. MonkeyLogic also synchronized behavioral events and gaze position. Gaze position was measured using infrared light-sensitive cameras, which visualized the eye through infrared-transparent mirrors (Qian and Brascamp, 2017), and commercially available eye tracking software (Eye Link II or SMI Research), which was converted to an analog signal that was sampled by MonkeyLogic/MATLAB (NIDAQ PCI-6229) at 1 kHz.

Animals viewed all stimuli through a custom-built mirror stereoscope (Carmel et al., 2010), such that images on the right side of the forward-positioned display were viewed by the right eye and images on the left by the left eye (the monitor was divided by a black, non-reflective septum). The total viewing distance from each eye to the monitor ranged between 46 and 57 cm, resulting in 20.5 to 34.5 pixels per degree of visual angle (dva). To facilitate fusion, an oval aperture was displayed at the edge of each half-screen. At the beginning of each experimental session, the stereoscope was calibrated via a behavioral task (Maier et al., 2008). The task required the animals to fixate the same location in visual space while being cued in one eye only. Gaze position was measured for each fixation location and compared across eyes. When gaze position was comparable for cuing in each eye, the mirrors were considered aligned.

### Behavioral Tasks

Both monkeys were trained to hold their gaze steady on a small (0.2 dva) spot presented in the center of the monitor for extended periods of time (2–3 s) while circular gratings or other stimuli were presented in the perifoveal visual field. If gaze remained fixed within 1 dva of the fixation cue for the duration of the trial, they received a juice reward. No other responses were required. These parameters apply to each the *Receptive Field*, *Tuning*, and *Dichoptic* paradigms described below.

### Neurophysiological Apparatus

Extracellular voltage fluctuations were recorded using acute linear multielectrode arrays (UProbe, Plexon Inc.; Vector Arrays, NeuroNexus) inside an electromagnetic radio frequency-shielded booth. The number of microelectrode contacts varied between probes (UProbe=24, Vector Arrays=32), but were always linearly spaced 0.1 mm apart. Voltage fluctuating signals were amplified, filtered, and digitized using a 128-channel Cerebus® Neural Signal Processing System (NSP; Blackrock Microsystems). Two signals were recorded and stored for subsequent offline analysis: A broadband (0.3 Hz – 7.5 kHz) signal sampled at 30kHz and a low frequency dominated signal (0.3 Hz – 500 Hz) sampled at 1 kHz. The low frequency dominated signal was used as a measure of the Local Field Potential (LFP). Spiking activity was extracted from the 30kHz broadband signal as described below.

The NSP also recorded non-neurophysiological analog signals related to monitor (i.e., a photodiode signal; OSI Optoelectronics) and eye position (i.e., voltage output of eye tracking system) which were digitized and stored at 30 and 1 kHz, respectively. The NSP stored timestamped event markers sent from the behavioral control system (MonkeyLogic). These timestamps and the photodiode signal were used to align the time-varying intracranial data with the occurrence of key visual and behavioral events.

### Experimental Design and Statistical Analysis

With the exception of LFPs (see above), all neurophysiological signals were extracted offline from the recorded broadband signal using custom-written code in MATLAB (2016a; The Mathworks, Inc.). We computed two measures of multi-unit activity, an *analog* signal (aMUA) and a *discretized* signal (dMUA). The analog signal was computed by high-pass filtering the broadband signal at 750 Hz with a 4^th^-order Butterworth filter and rectifying (Supèr & Roelfsema, 2005). The discretized signal was computed by applying a time-varying threshold to the envelope of the broadband signal, with an impulse recorded at every time point where the signal envelope exceeded the threshold. This procedure is analogous to the initial step of spike sorting without an ensuing step of cluster analysis. For both the threshold and the envelope computations, we began by lowpass-filtering the 30 kHz-sampled voltage signals at 5 kHz with a second order Butterworth filter, downsampling by a factor of 3, high pass-filtering at 1 kHz with a second order Butterworth filter, and then rectifying the resulting data. For the envelope computation, we then downsampled the signal by a factor of 3. For the threshold computation, we smoothed the signal by convolving the data with a 1 s boxcar function and then multiplied the result by 2.2. To recover temporal information, we extracted +/- ms of data from the original signal for each time point where the envelope exceeded the threshold. We then adjusted the time point to correspond with the point of maximum slope within this window, i.e. aligning to the spike waveform. For all MUA signals, a “multiunit” describes the neuronal signal extracted using the techniques described above from a single microelectrode from a single penetration.

We also extracted single-unit activity (SUA) via Kilosort, an unsupervised machine-learning spike-sorting algorithm (Pachitariu et al., 2016). A major benefit of using Kilosort for extracting single-unit activity from linear electrode arrays is that neurons that produce a signal on more than one microelectrode are not counted more than once. We used the default parameters for sorting and cluster merging, so we focus here on our customized post-processing steps only. For all clusters detected by Kilosort, we extracted +/- 1 ms of data around each spike from the original broadband signal for each simultaneously recorded electrode contact. We averaged across all extracted impulses to create a spatiotemporal map of the spike waveform (time x electrode contacts). To be included in the study, the region of the spatiotemporal waveform map that exceeded +/- 30% of maximum modulus had to span less than 0.9 ms and 3 neighboring microelectrode channels (0.3 mm). Clusters that met these criteria were localized to the microelectrode with the largest amplitude.

Spike rates for each dMUA and SUA were limited to 1 kHz (except **Figure 2f**). In all cases, unit data was convolved using a Poisson distribution resembling a postsynaptic potential (Hanes et al., 1995), with the spike rate (*R*) computed at time (*t*):

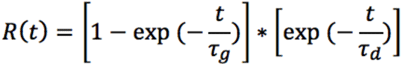

where *τ_g_* and *τ_d_* are the time constants for growth and decay, respectively. Data from previous studies suggest values of 1 and 20 for *τ_g_* and *τ_d_* respectively (Sayer et al., 1990). For aMUA, this kernel was convolved with the filtered and rectified time-varying signal directly. For dMUA and SUA, impulse times were converted to a time-varying signal using 0 to represent time points without an impulse and 1 for time points where an impulse was detected, and then convolved. After convolution, the signal was multiplied by the sampling frequency to convert units to spikes per s.

Current source density (CSD) analysis was performed on the LFP signal using an estimate of the second spatial derivative appropriate for multiple contact points (Nicholson and Freeman, 1975).

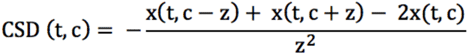

where *x* is the extracellular voltage recorded in Volts at time *t* from an electrode contact at position *c*, and *z* is the electrode inter-contact distance (0.1mm). In order to yield CSD in units of current per unit volume, the resulting CSD from the formula above was multiplied by 0.4 S/mm as an estimate of cortical conductivity (Logothetis et al., 2007).

### Receptive Field Paradigm and Analysis

Subjects fixated a central cue while circular patches of static random noise were displayed at pseudorandomized locations within a pre-determined virtual grid (**Figure 1a**). Noise patches were generated by multiplying a 2D-Gaussian with a randomized patch of full-contrast black and white pixels and then cropping at radius = 3*σ. Stimulus size, grid spread, and grid step were manually determined by the experimenter each session, and typically included both a coarse mapping phase with large stimuli (σ = 1–2 dva) and grid spacing (i.e., 2–5 dva) covering a whole visual quadrant (theta = 0–90°, eccentricity = 0–8 dva) and a finer mapping phase (σ = 0.125–0.5 dva; spacing = 0.25–1 dva) focused on the portion of the visual field thought that contained the receptive fields of the neurons under study. Up to 5 stimuli were shown per trial, for 200 ms each with 200 ms blank periods interleaved. We used retinotopic 3D Receptive Field Matrices (RFMs) (Cox et al., 2013) to compute spatial maps of neuronal responses as a function of visual space (**Figure 1b-c**). For each stimulus presentation, the response of a given unit (i.e. MUA from a single microelectrode contact or derived SUA) was averaged to produce a single scalar value (0.05–0.25 s from stimulus onset). These scalar values were converted to units of z-score (n = all stimulus presentations, regardless of stimulus parameters). Then, for each trial, the retinotopic portion of the RFM corresponding to the stimulus location (radius = Gaussian σ) was filled with that value producing a matrix consisting of one dimension for vertical visual space, one for horizontal visual space, and one for stimulus presentations. This third dimension was then collapsed via averaging, producing a spatial map of the unit’s response. Receptive field centers and extents were computed by fitting an oval to the largest, contiguous patch of the spatial map that exceeded 1 z-score (**Figure 1c-e**).

**Figure 1:**
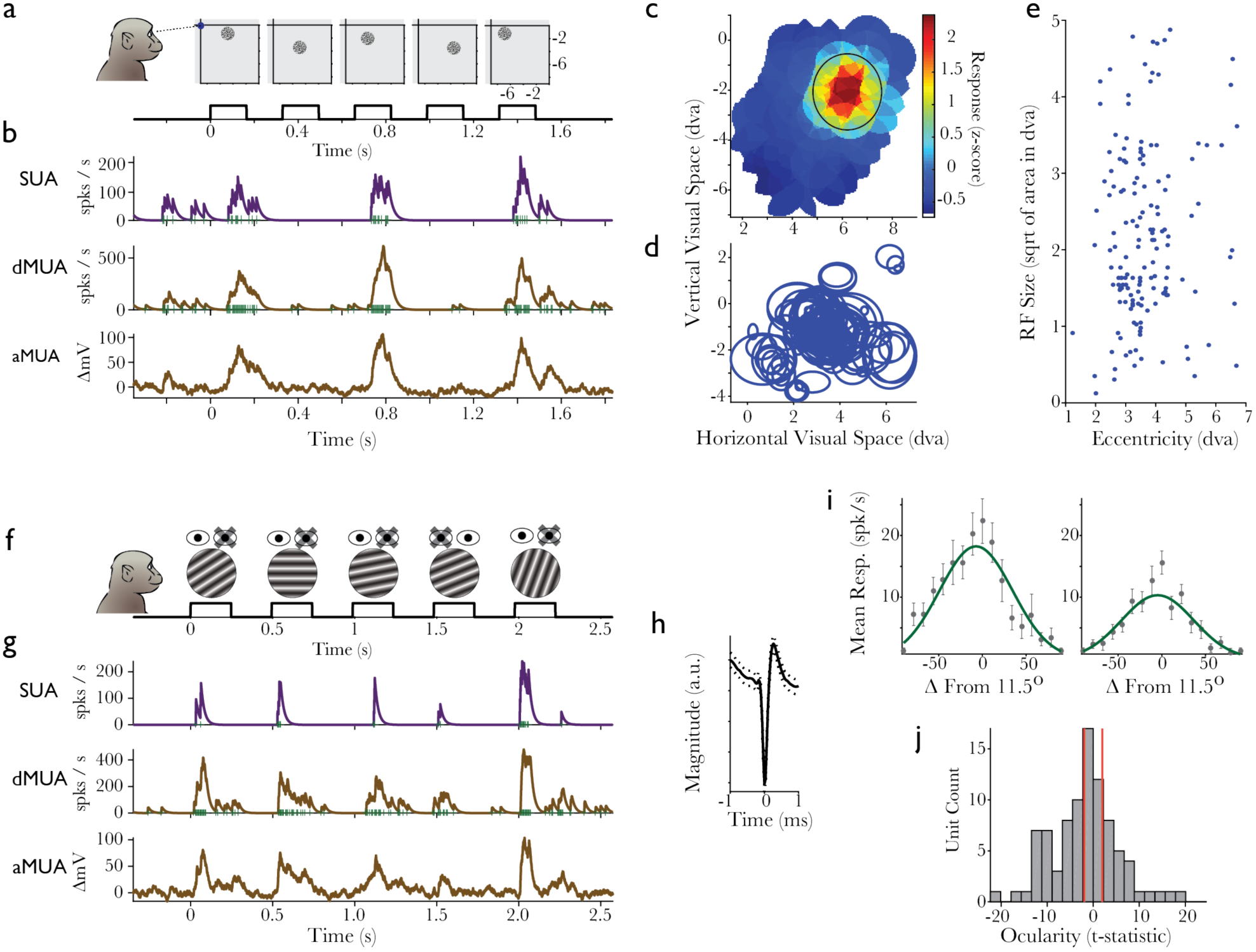
Determining units’ receptive fields and tuning. (a) Receptive field mapping using static random noise patches. (b) Single-trial activity of each SUA, dMUA, and aMUA evoked by the stimuli in a. For SUA and dMUA, green vertical lines indicate a detected spike and blue lines are the result of convolving spikes with a PSP kernel. For aMUA, the blue line is the result of convolving the rectified, high-passed extracellular voltage signal with the same kernel. (c) Example SUA Receptive Field Matrix (RFM) showing neuronal responses as a function of visual space. Z-axis is in units of z-score across all stimulus presentations (n = 658). Black circle signifies the receptive field boundary. (d) RF boundaries for all SUA (n = 152), x-axis in rectified units. (e) RF area as a function of eccentricity (f) Tuning Paradigm. (g) Single-trial activity evoked by the stimuli in f, convention as in b. (h) Mean waveform for SUA shown in g and i. Dotted line is SEM. (i) SUA as a function of grating orientation for presentations to each the contralateral (left) and ipsilateral (right) eye. Error bars are SEM. Green lines are Gaussian fits to data. Given the preference of this neuron, 0 was set to an actual on-screen orientation of 11.5-degree counterclockwise tilt from horizontal (j) Histogram of ocularity indices, computed as a t-statistic between monocular responses, across all units showing a significant main effect of orientation (n = 95). Red lines approximate the critical t-value required for a unit to be classified as either equiocular (middle of histogram) or ocular-biased (tails of histogram).

### Tuning Paradigm and Analysis

Subjects fixated a central cue while circular grating patches appeared at a single location in parafoveal visual space, determined by the results of the previous *Receptive Field Paradigm and Analysis*. Up to five stimuli were shown per trial, typically for 250 ms each with 250 ms blank periods interleaved. Each sinusoidal grating patch could vary in eye, orientation, spatial frequency, and phase. Binocular stimulation always consisted of identical stimuli between the eyes (dioptic). Parameters were determined by the experimenter each session based on their assessment of audible multiunit responses to the stimuli described here, and typically included some combination of monocular and dioptic stimuli varying in orientation by at least five steps between 0° and 180°. Contrast was held steady at a Michelson contrast of 0.9 or 1.0 (all contrasts values in this study are reported in Michelson contrast). Stimulus-evoked responses were averaged 0.05–0.25 s from stimulus onset for each trial. When computing orientation tuning curves (**Figure 1i**) as well as when determining ocularity (**Figure 1j**), all non-relevant parameters were matched (e.g., for comparison between responses evoked by each eye, orientation and phase were balanced between the eyes). Preferred orientation was typically consistent across units collected on different V1-residing multielectrode in a single penetration of the linear array (see *Intersession Alignment*; a detailed description of the laminar distribution of orientation tuning is the focus of upcoming work), and the orientation tuning of units included in this study (see *Determining Orientation Tuning and Ocular-Bias*) corresponded to the stimulus orientations used in the *Dichoptic Paradigm*.

### Dichoptic Paradigm and Analysis

Subjects fixated a central cue while circular grating patches appeared at a single location in parafoveal visual space, determined by the results of the previous *Receptive Field Paradigm and Analysis*. Up to three stimuli were shown per trial, typically for 500 ms each with 500 ms blank periods interleaved. Sinusoidal grating patches varied between two orientations — determined by the results of the previous *Tuning Paradigm and Analysis* — and contrast (3+ steps between 0.1 and 1.0 Michelson contrast) with spatial frequency and phase held constant. Stimuli were shown to each eye in isolation (monocular, M) and to both eyes simultaneously such that in some presentations the same orientation was shown to both eyes (binocular congruent, BC) and on other trials the orientation differed between the eyes (binocular incongruent, BI). As the central aim of the study was to examine the effects of non-dominant eye stimulation on neuronal responses to dominant eye stimulation with a preferred stimulus, all stimulus conditions were defined relative to each neuron’s preferred monocular condition. On the majority of penetrations (N = 22 of 30), this design was fully balanced, however a few penetrations contained only the binocular congruent or binocular incongruent conditions (**Table 1**). When computing binocular modulation, both via the t-statistic and Michelson contrast, the monocular condition always served as the subtrahend (e.g., BC - M or BI - M), such that positive values represent greater response for the binocular conditions and negative values represent greater response for the monocular condition.

**Table 1:** Single unit distribution.

|  | Total Penetrations | Total Single Units | Orientation Selective | Orientation Selective and Ocular-Biased | Orientation Selective and Equiocular |
| --- | --- | --- | --- | --- | --- |
| *Congruent* | 24 | 115 | 78 | 57 | 32 |
| *Incongruent* | 29 | 135 | 85 | 66 | 28 |
| *Congruent OR Incongruent* | 30 | 152 | 95 | 70 | 37 |
| *Congruent AND Incongruent* | 22 | 98 | 68 | 53 | 23 |

Total number of single units isolated across 30 total penetrations (6 in Monkey 2), for each population (columns) and binocular stimulus condition (rows). See Methods for details on single-unit extraction and classification.

### Determining Orientation Tuning and Ocular-Bias

As stimulus orientation is the critical differentiating variable in the *Dichoptic Paradigm*, we limited our analysis to orientation-tuned units. Orientation tuning was determined by analyzing monocular and dioptic data from each the *Tuning* and *Dichoptic* tasks and testing for a significant main effect of orientation via an ANOVA. All other stimulus parameters (orientation, phase, contrast, spatial frequency, monocular eye v. dioptic) were included as groups in the ANOVA. Reponses were averaged over a time period ranging 0.05–0.25 s from stimulus onset. Ocularity of each unit was determined similarly. However, as there are only ever 2 monocular conditions, we performed a t-test between eyes after balancing all other parameters (see also *Tuning Paradigm and Analysis*). Responses for the ipsilateral eye served at the minuend, such that positive t-statistics represent greater responses for the ipsilateral eye and negative t-statistics represent greater responses for the contralateral eye (**Figure 1j**). If a unit exhibited a significant response difference between stimulation in each eye, we classified it as ocular-biased. Otherwise, it was considered equiocular.

### Intersession Alignment

The relative depth of each microelectrode in cortex was determined using several neurophysiological criteria: The upper and lower bounds of V1 were determined algorithmically using data from the *Receptive Field Paradigm* and the *Tuning Paradigm*. First, we used the magnitude of the stimulus-evoked dMUA response to a grating stimulus placed over the RF to determine which microelectrodes gave a reliable stimulus-evoked response (paired-sample t-test of mean response for the time periods −50–0 ms from stimulus onset and 50–100 ms, FWER = 0.05; using all stimulus presentations regardless of parameters). We also extracted receptive fields from dMUA as described in *Receptive Field Paradigm and Analysis*. As dMUA does not rely on single-unit isolation, we were able to compute RFMs for each microelectrode contact (**Figure 2a**) and use the 1 z-score criteria to determine the location and extent of each receptive field. An aggregate receptive field center for each cortical location was calculated by all concurrently measured receptive fields averaging across depth. Then, we checked that all dMUA-derived RFMs had a center within 0.25 dva of the aggregate receptive field center. In this way, receptive field mapping information was used to identify spatial bounds of V1. Typically, this RFM-based analysis eliminated superficial channels for not reaching the 1 z-score criteria anywhere within their RFM and deep microelectrodes either due to either a deviation in mapped center (possibly due to penetrating another fold of cortex) or for not reaching the 1 z-score criteria anywhere within their RFM. In cases were the V1 bounds extracted from the evoked response differed from those extracted from the RFMs, we averaged the two sets of measures, rounding inwards.

**Figure 2:**
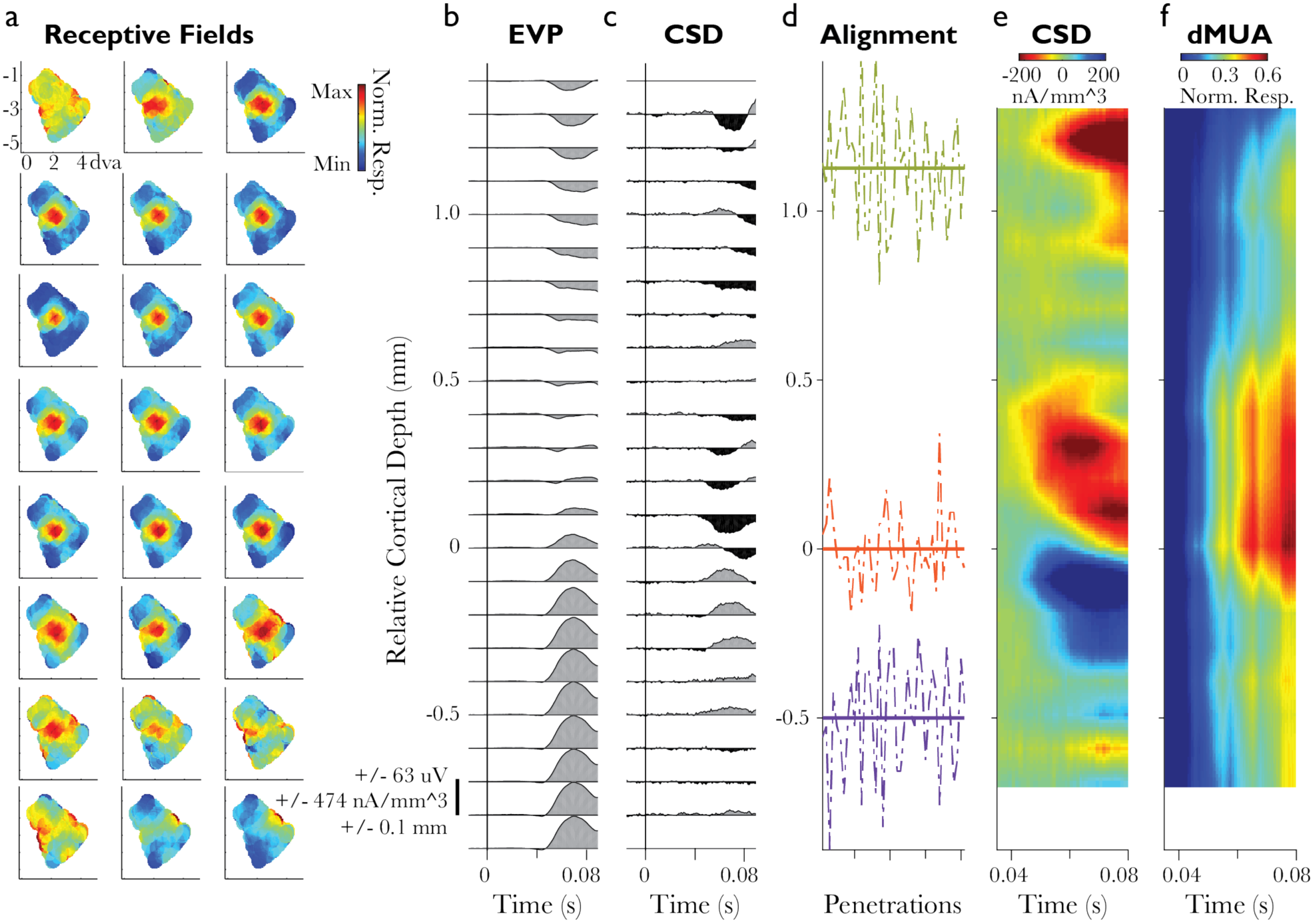
Localizing units to cortical depth. (a) RFMs extracted from the dMUA signal recorded on each microelectrode in an example V1 linear array penetration. Top left panel is the most superficial microelectrode, bottom right panel is the deepest. Conventions for each RFM plot are as in Figure 1c, though here the z-axis is scaled individually for each microelectrode. (b) Evoked LFP following a brief (88 ms) black-to-white flash of the display monitor (n = 271 trials). (c) Evoked CSD extracted from data in b. Negative deflections (black) indicate current sinks and positive deflections (gray) indicate current sources. (d) Functionally-determined depth of each the top of V1 (green), bottom of V1’s middle granular input (orange), and the bottom of V1 (purple) across all V1 penetrations (abscissa) examined for this study (n = 41, 30 of which contained both SUA and the appropriate stimulus conditions for the main study; ordinate in relative units). On average, the bottom of Layer 4 was 1.2 mm from the top of V1 and 0.5 mm from the bottom of V1. (e) Flash-evoked CSD, averaged across penetrations and interpolated in the depth dimension only (10 pts per 0.1 mm^3). (f) Flash-evoked dMUA, normalized to the maximum response and then averaged across penetrations.

We used a well-established and histologically-verified neurophysiological method to functionally determine the location of V1’s granular input layer (layer 4C; **Figure 2b-c**). Specifically, CSD analysis of visual responses to brief visual stimulation has been shown to reliably indicate the location of the primary geniculate input to V1 in the form of a distinct current sink that is thought to reflect combined excitatory post-synaptic potentials of the initial retinogeniculate volley of activation (Mitzdorf, 1985; Maier et al., 2010; 2011; Dougherty et al., 2015; Cox et al., 2017). For each penetration with the laminar multielectrode array, CSD analysis was used to resolve this prominent initial current sink immediately following stimulus onset. The bottom of this sink was used as a marker of the transition between granular layer 4C and the deeper layer 5. Thus, using a combination of criteria, each penetration was assigned three reference microelectrodes representing each the top of V1, Layer 4C/5 boundary, and bottom of V1 (**Figure 2d**). These points were used to align and average data across electrode penetrations and recording sessions, resulting in 0.1 mm +/- 0.05 mm resolution across V1’s cortical depth (**Figure 2e-f**).

## Results

The first objective of this study was to compare the time course of V1 spiking between monocular stimulation and binocular stimulation. A mirror stereoscope was used to display stimuli independently to each eye while keeping the overall level of illumination constant. The spiking activity of V1 neurons was assessed via intracranial recordings with linear multielectrode arrays.

For each electrode penetration, the receptive field location and tuning properties of the recorded neurons were characterized using a series of visual stimuli that were progressively fine-tuned by the experimenter (**Figure 1**, Methods). The goal of this initial phase of the experiment was to identify the preferred orientation and ocular preference of the recorded V1 neurons so that the parameters of the main experiment (**Figure 3a-c**) could be customized to the neurons’ response preferences. Following the definition of dCOS used by previous studies (Sengpiel et al., 1998), three main stimulus conditions were defined relative to each neuron’s preferred monocular condition:

<u>Monocular (M)</u>: stimulus in the neurons’ preferred (dominant) eye at the neurons’ preferred orientation; no grating stimulus in non-dominant eye (contrast = 0). In plots, shown in *black*.
<u>Binocular Congruent (BC)</u>: dominant eye stimulus set to the neurons’ preferred orientation; non-dominant eye stimulus set to the neurons’ preferred orientation. Contrast in non-dominant eye ~2x that of the dominant eye. In plots, shown in *blue*.
<u>Binocular Incongruent (BI)</u>: dominant eye stimulus set to the neurons’ preferred orientation; non-dominant eye stimulus oriented orthogonally to the neurons’ preferred orientation. Contrast in non-dominant eye ~2x of the dominant eye. In plots, shown in *red*.

**Figure 3.**
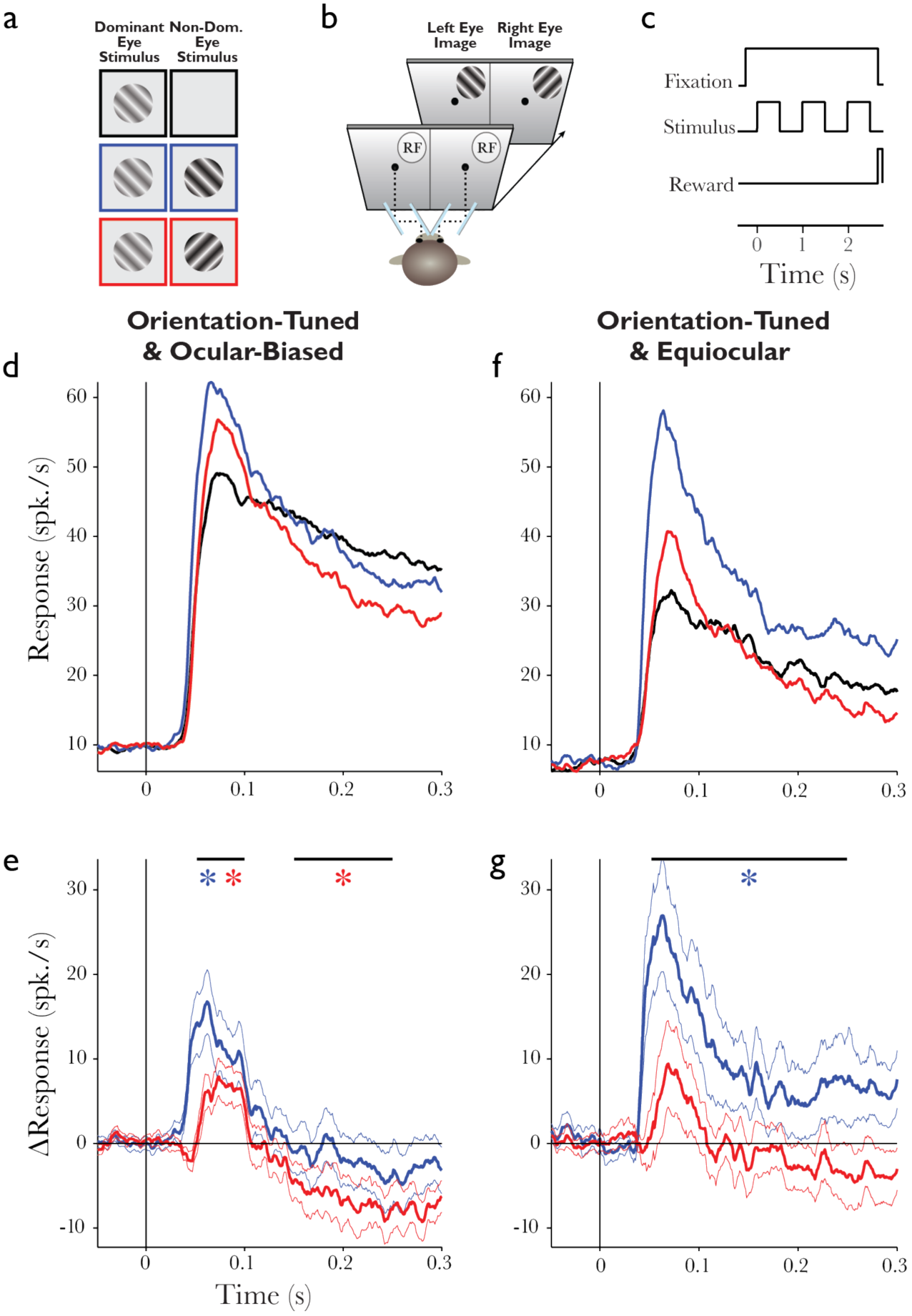
Neuronal responses to binocular stimuli. (a) Main stimulus conditions in the Dichoptic Paradigm. M = Monocular, BC = Binocular Congruent BI = Binocular Incongruent. Colored outline serves and legend for subsequent plots. Note that the stimulus in the non-dominant eye was always set to 100% contrast. (b) Subjects viewed a single divided display through a custom-built mirror stereoscope. (c) Stimuli appeared on the display for 0.5 s with a 0.5 s interstimulus interval. (d) SUA evoked by each of the main dichoptic stimulus conditions for ocular-biased units. Stimulus onset occurs at 0 s, mean across units (n = 53). (e) Binocular-evoked SUA minus monocular-evoked SUA for each the congruent (blue) and the incongruent (red) binocular conditions for ocular-biased units. Subtraction taken for each unit and then averaged across populations (thin line = SEM). (f-g) Same as d and e, but for equiocular units (n = 23).

We focused our study on the spiking activity of individually isolated V1 neurons, a.k.a. single-unit activity (SUA, n = 152; see Methods). As stimulus orientation was the critical differentiating feature between congruent and incongruent binocular stimulation conditions, we limited our analysis to V1 units with a significant main effect of stimulus orientation (n = 95). We further separated these units based on ocularity (**Table 1**; **Figure 1j**). Specifically, we divided units that exhibited significant response preferences for one eye, which we termed *ocular-biased*, from units that did not, which we termed *equiocular*. All equiocular units showed a slight but insignificant difference in mean firing between the eyes, which allowed us to nonetheless determine a preferred (dominant) and non-preferred (non-dominant) eye.

### Temporal dynamics of binocular V1 responses

First, we examined stimulus-evoked spiking responses for ocular-biased units (**Figure 3d&e**; *black* = monocular, *blue* = binocular congruent, *red* = binocular incongruent). We found that initial activity evoked by both types of binocular stimuli (congruent and incongruent) significantly exceeded that of the monocular stimulus (congruent vs. monocular, *t*(52) = 3.81, *p* < .001; incongruent vs. monocular *t*(52) = 3.74, *p* < .001), with the congruent evoking an even greater response than the incongruent stimulus. However, this binocular enhancement of V1 responses was limited to the transient phase of the visually-evoked response (~100 ms). During the sustained response period following the initial transient, binocular responses decreased to the point where binocular responses, both to the congruent and the incongruent stimuli, were reduced relative to monocular stimulation, with the incongruent reduction being both larger and more reliable across the population (congruent vs. monocular, *t*(52) = −0.66, *p* = 0.510; incongruent vs. monocular *t*(52) = −2.51, *p* = .015). Applying the same analysis to equiocular units revealed a similar temporal response pattern for the transient response but not for the sustained response. Specifically, sustained responses for the congruent condition were greater than sustained responses to monocular stimulation (**Figure 3f&g**, *blue* vs. *black*; *t*(22) = 3.95, *p* < .001). This pattern is indicative of orientation-specific binocular enhancement exclusive to equiocular units.

To statistically assess the reliability of these findings, we separately compared each binocular condition to the monocular condition across both the initial transient and the sustained period of V1 visual responses. Specifically, we defined two temporal analysis windows: an <u>early window</u>, 50–100 ms following stimulus onset, which corresponds to the *initial transient* of the response, and a <u>late window</u>, 150–250 ms following stimulus onset, which corresponds to the *sustained* part of V1’s visual response. In the early time window, a large fraction of ocular-biased neurons showed response enhancements for binocular stimulation. Specifically, 36% of ocular-biased neurons exhibited a significantly greater response for the binocular congruent condition compared to the monocular condition and 25% for binocular incongruent compared to monocular (Student’s t-test, α = 0.05, two-tailed). In the later, sustained response window, the proportion of ocular-biased units exhibiting greater responses for the binocular conditions decreased, and the proportion of units demonstrating reduced responses for binocular conditions increased such that 25% of units exhibited a significantly reduced response for the binocular incongruent condition compared to the monocular condition and 13% for binocular congruent compared to monocular. Equiocular units exhibited a similar pattern of responses across both the early and late windows for incongruent compared to monocular stimulation, with 22% of units exhibiting significantly greater binocular activity in the early window and 22% of units exhibiting significantly lesser binocular activity in the late window. However, this pattern differed when stimuli were congruent between the two eyes, with 48% of equiocular neurons exhibiting significantly greater binocular activity in the early window and 13% in the later window. These population data confirm the two epochs of binocular modulation in V1 observed in the spiking time courses: An early period where response for binocular stimuli are increased relative to monocular stimulation followed by rapid response reduction for the majority of V1 neurons. The only exception to this interocular suppression was observed among the population of equiocular V1 neurons that do not encode eye-specific information. Even more interestingly, this exception was contingent upon – and thus indicative of - congruent stimulation between the eyes.

To further quantify the effects of binocular stimulation across our sample of V1 neurons, we computed a normalized binocular modulation index in the form of a Michelson Contrast for each binocular stimulation condition compared to monocular stimulation (Binocular Response – Monocular Response/ Binocular Response + Monocular Response). This analysis allowed us to quantitatively investigate the distribution of binocular modulation across the entire neuronal population as well as to examine the degree to which V1 neurons distinguish between congruent and incongruent binocular stimulation. Specifically, the correlation between binocular congruent and incongruent modulation is indicative of the degree to which interocular orientation differences impact binocular responses. A strong positive correlation between binocular congruent and incongruent responses indicates that the V1 binocular population response is non-specific to stimulus orientation mismatches between the eyes. Alternately, if differences in orientation between the eyes impact binocular response modulation (e.g., causing suppression or enhancement), we expect responses for the binocular 398 stimulation conditions to be either uncorrelated or negatively correlated. We found that for both ocular-biased and equiocular neurons, modulation was strongly positively correlated throughout the initial transient captured by the early response window (**Figure 4a-b**, respectively: *r*(51) = 0.45, *p* < 0.001; *r*(21) = 0.40, *p* = 0.059). This result suggests that the transient binocular response enhancement in V1 following onset of visual stimulation is indifferent to whether or not the images in the two eyes are congruent. However, in the later, sustained part of the response, correlations generally decreased (**Figure 4c-d**, respectively: *r*(51) = 0.20, *p* = 0.15; *r*(21) = 0.24, *p* = 0.280), suggesting that binocular congruency is computed in a secondary step.

**Figure 4:**
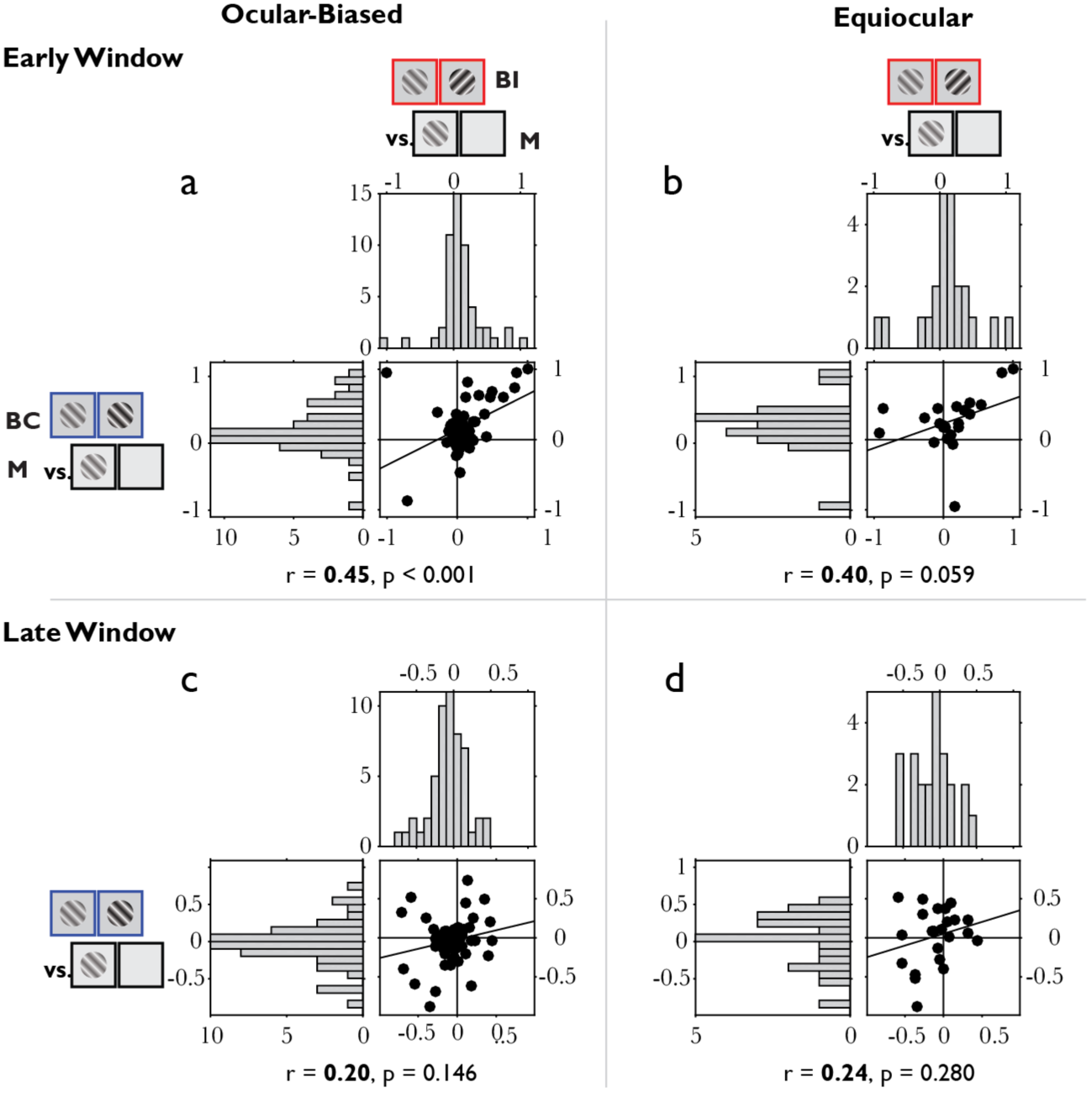
Correlation of V1 binocular modulation in each the early, transient response and the late, sustained response. Binocular modulation for each the early (a,b) and late (c,d) part of the visual response across both ocular-biased units (a,c; n = 53 units) and equiocular units (b,d; n = 23 units). In each panel, the top horizontally-oriented histogram indicates the distribution of incongruent binocular modulation (BI vs. M) and the left vertically-oriented histogram indicates the distribution of congruent binocular modulation (BC vs. M). The central scatterplot contains the same data as each histogram, with modulation for the congruent condition on the ordinate and modulation for the incongruent condition the abscissa. The coefficient (r) and p-value for each correlation is listed below the corresponding scatter plot.

### Laminar profile of V1 binocular response modulation

We performed all V1 electrode penetrations using linear microelectrode arrays. As a result, we were able to identify the upper and lower bounds of V1 of each penetration as well as the bottom of layer 4C using a variety of electrophysiological criteria (**Figure 2**, Methods). Using these functionally defined cortical depth markers, we assigned each microelectrode contact of the linear array, and the corresponding units recorded on those contacts, a laminar location relative to the bottom of layer 4C. Then, we examined the distribution of the binocular modulation index as a function of laminar location. Comparing the binocular incongruent stimulation with stimulation of one eye alone, we observed that the transient (early-window) binocular enhancement was expressed predominantly in the middle and superficial layers (0.0–1.2 mm above the L4C/5 boundary) and not in the deep layers of V1 (−0.7–0.0 mm below the L4C/5 boundary; **Figure 5a**). During the sustained response (late window), binocular modulation decreased in both the middle and superficial layers. However, while neurons in the superficial layers (0.5–1.2 mm above L4C/5) exhibited strong response reduction relative to monocular stimulation (M), neurons in the middle layers (0.0–0.4 mm above L4C/5) showed minimal differences between binocular incongruent and monocular stimulation (**Figure 5b**).

**Figure 5:**
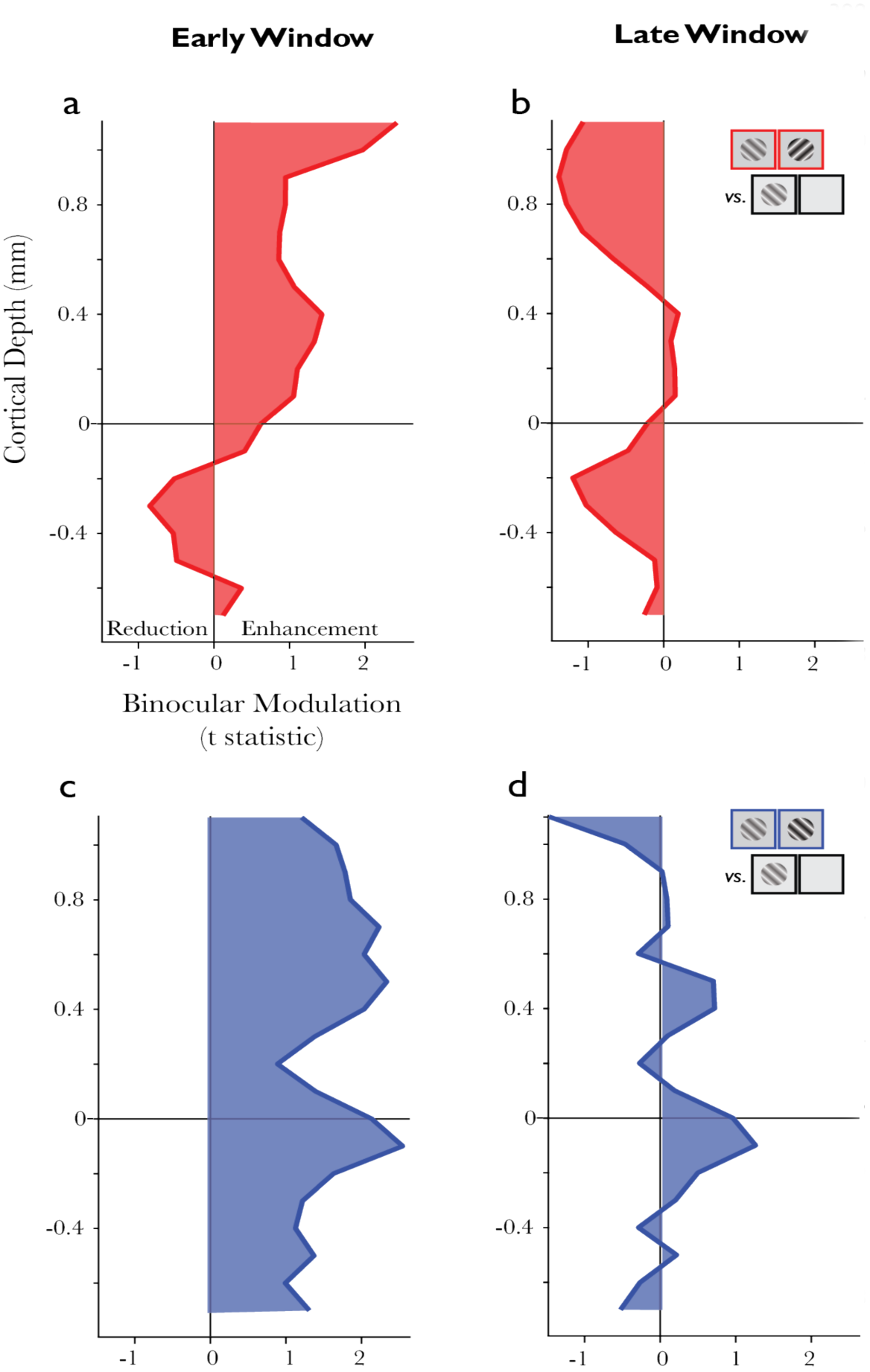
Binocular Modulation across VI’s Layers. (a) Pooled incongruent binocular modulation across layers in units of t-statistic for the early analysis window corresponding to the transient response. To compensate for low sampling in some layers, the mean at a given depth was calculated by pooling units across one depth unit above and below. (b) Same as in a except analysis is across the late window corresponding to the sustained response. (c) Pooled congruent binocular modulation, conventions as in a. (d) Same as in c except analysis is across the late window corresponding to the sustained response.

Repeating the same analysis for the binocular congruent stimulation condition revealed a different laminar pattern of activity. Initially, all layers exhibited binocular enhancement in the early, transient response window (**Figure 5c**), with the lower portion of the middle, granular layers (0.0–0.3 mm above L4C/5) showing the least binocular enhancement. Then, during the late, sustained response, binocular modulation decreased, with neurons across most cortical depths showing minimal response differences between binocular and monocular stimulation (**Figure 5d**). However, on an individual basis, units exhibiting sustained binocular response enhancement clustered predominately in the superficial layers (0.4–0.6 mm above L4C/5) as well as just below the granular layer (−0.3–0.0 mm below L4C/5).

## Discussion

We found that the temporal dynamics of V1 responses to each congruent and incongruent binocular stimulation are indicative of a series of computational steps that the visual system employs in processing the concordant and discordant views of the two eyes. Both V1 neurons that were significantly selective for one eye as well as V1 neurons that were binocularly driven initially responded more vigorously when both eyes were stimulated, suggesting some form of (sub-additive) binocular summation. About 150 ms following this initial binocular response facilitation, both groups of neurons exhibited a relative decrease in response magnitude compared to monocular stimulation, signifying some form of interocular suppression. This widespread V1 response suppression only occurred if the stimuli in the two eyes were of orthogonal orientation (incongruent), posing significant interocular conflict. Neurons with significant response preference for one eye over the other also underwent interocular suppression when the stimuli in the two eyes were of the same orientation (congruent) and thus perceptually fusible. However, binocularly driven neurons that lacked significant ocular preference (equiocular neurons) showed the opposite pattern in the form of sustained binocular summation when both eyes were stimulated rather than one. Taken together, these results suggest that following an initial boost in firing caused by stimulation of both eyes rather than one, V1 neurons rapidly differentiate between congruent and incongruent stimuli across the two eyes, resulting in widespread reduction of visual responses when interocular conflict arises. This transition from response enhancement in the transient response to response reduction in the sustained response may serve as initiator for secondary mechanisms of interocular conflict resolution resulting in stereopsis, diplopia or binocular rivalry.

### Response dynamics in V1

We still know little about how cortical areas are organized in the temporal domain (Murray et al., 2014). Temporal response characteristics of V1 neurons have been studied extensively in the context of orientation tuning (Celebrini et al., 1993; Ringach et al., 1997), where inhibition rapidly follows excitation to increase orientation selectivity (Ringach et al., 2003; Shapley et al., 2003; Litwin-Kumar et al., 2016; but see: Mazer et al., 2002). For the neuronal response dynamics reported here, response reduction for binocular stimuli emerges within ~50 ms of the initial transient. One interpretation is that this initial response, where activity for binocular stimulation exceeds the magnitude of monocular stimulation, is driven predominately by feedforward excitation while the subsequent response reduction for binocular stimulation is a result of cortical-cortical inhibition. This inhibition could be mediated by short-range horizontal connections between neighboring ocular dominance columns. While long-range (>0.5mm) horizontal connections in the superficial layers of primate V1 tend to be homophillic for ocularity and orientation, short-range (<0.5mm) horizontal connections are dense and more promiscuous (Malach et al., 1993; Yoshioka et al., 1996; Das and Gilbert, 1999; Stettler et al., 2002; Tychsen et al., 2004). However, the observation that congruent binocular stimuli evoke response enhancement in V1 equiocular cells, suggests that interocular inhibition is bypassed or superseded in this condition. The functional differences between equiocular and ocular-biased neurons suggest that each group receives different input from the two eyes, with equiocular neurons receiving nearly-equivalent excitatory drive from each of the two eyes. It thus seems conceivable that the latter group of neurons features unique reciprocal excitatory connections between cells that share orientation preferences.

### Relation to previous dCOS literature

The comparisons between the binocular incongruent and monocular stimulation conditions in this study are comparable to the stimulation conditions of the previously described phenomenon of dichoptic cross-orientation suppression (dCOS). The vast majority of work on dCOS has been carried out in area 17 of anesthetized, paralyzed cats (Sengpiel and Blakemore, 1994; Sengpiel et al., 1998; Walker et al., 1998; Sengpiel, 2005). While the anatomical organization of binocular vision differs significantly between cats and primates (Wilson and Cragg, 1967; Hendrickson et al., 1978; Yoshioka et al., 1996; Heesy et al., 2011, Dougherty et al., 2018), area 17 in the cat and V1 in primates are generally considered functionally homologous. One goal of this study was to examine the degree to which previous dCOS studies in area 17 can be generalized to the awake primate. To a first approximation, our data reveal that dCOS occurs in primates to a similar degree as in cats. Previous studies have suggested that dCOS is mediated by intra-cortical inhibition (Sengpiel, 2005). This conclusion was based on the observation that dCOS exhibits differential contrast-gain control compared to monocular cross-orientation suppression, dCOS’ sensitivity to visual adaptation and the rate of stimulus motion, and the ability to extinguish dCOS with GABA-antagonist in area 17. Our finding that units outside V1’s main retinogeniculate input layer exhibit the strongest orientation-specific binocular suppression corroborates this view that dCOS arises from intra-cortical interactions. Our findings in V1 also agree with cat studies on the frequency of dCOS across neurons, with over 50% of our ocular-biased neurons exhibiting suppression after ~150 ms, as well as the magnitude of suppression, which hovered around ∆10 spikes / sec (Sengpiel et al., 1998). Another study in anesthetized macaques reported similar percentages for adult subjects (Endo et al., 2000), providing further support that application of anesthetics does not significantly alter this fundamental mechanism.

### Relation to binocular rivalry

Another popular approach to studying binocular integration on the level of single neurons in visual cortex have been binocular rivalry paradigms (see Logothetis, 1998; Leopold, 2012; Schmid and Maier, 2015 for review). In fact, most models of interocular suppression are designed to specifically account for perceptual alterations in binocular rivalry (Laing and Chow, 2002; Wilson, 2003; Grossberg et al., 2008), and generally postulate competition between neurons representing each eye, mediated by reciprocal inhibition between eye-specific neuronal populations (Tong et al., 2006; Blake and Wilson, 2011; Brascamp et al., 2013).

However, neuronal correlates of binocular rivalry are defined differently than dCOS. In binocular rivalry, neuronal responses are studied relative to the subject’s perceptual experience (Leopold and Logothetis, 1996; Sheinberg and Logothetis, 1997; Polonsky et al., 2000; Gail et al., 2004; Macknik and Martinez-Conde, 2004; Wunderlich et al., 2005; Maier et al., 2007; 2008; Wilke et al., 2009; Keliris et al., 2010; Maier et al., 2012; Bahmani et al., 2014; Xu et al., 2016). That is, the neuronal signals are typically investigated in order to identify populations of neurons that correlate with the fluctuating percept during interocular conflict. The general assumption of these studies is that the population of neurons representing the non-perceived stimulus are suppressed. In contrast, dCOS is defined from a neuron’s perspective, such that all comparisons are to that neuron’s preferred stimulus and dominant eye, and thus agnostic to perception (Sengpiel and Blakemore, 1994; Sengpiel et al., 1995b; Endo et al., 2000; Sengpiel, 2005). Across V1, there are neurons that prefer each eye-orientation combination. Thus, for any given binocularly incongruent stimulus configuration, dCOS will cause widespread response reduction among all neurons representing both eyes as well as both rivaling stimuli. If sensory drive from the two eyes is balanced between the eyes — as is characteristic for normal, unaltered vision (see (Wolfe, 1984; Wilke et al., 2003; Klink et al., 2010; Yang et al., 2015) for interesting exceptions) — then dCOS does not favor either eye or stimulus at the population level and instead remains constant while perception fluctuates between the two eyes’ views. This invariance supports the supposition that dCOS is a fundamental neuronal computation that is not linked directly to the perceptual outcome of ongoing binocular rivalry alternations.

dCOS may serve as a signaling mechanism of binocular conflict that precedes and ultimately paves the way for binocular rivalry alternations (Sengpiel et al., 1995a; 1998; Baker et al., 2007). In this context, it is noteworthy that perceptual dominance of one stimulus over another is not instantaneous in binocular rivalry. Subjects require at least 150 ms of exposure to orthogonal dichoptic gratings before one eye’s view is selected at the expense of the other (Wolfe, 1983). The timing of this perceptual property of binocular rivalry is well matched to the two distinct phases of binocular integration reported here. Specifically, it seems that the initiation of response reduction for binocular incongruent stimuli is coincident with the initiation of perceptual alternation in binocular rivalry, lending some support for the hypothesis that the onset of dCOS may be related to the initiation of binocular rivalry.

### Implications for binocular processing

The evolution of binocular signals across the V1 microcircuit is particularly informative regarding the sequence of computational processes underlying integration of the two eyes’ signals, especially in view of the classic canonical microcircuit model. According to this model, visual information flows from the granular input layer (layer 4, or layer 4C in primates) to superficial layers (layers 2/3) and then to the deep layers (layers 5/6; Callaway, 1998; Douglas and Martin, 2004). We found that neurons in both the middle granular layer and the superficial layers of V1 exhibit early enhancement for incongruent binocular stimulation, while neurons in the deep layers did not. In the later response period, granular layer activity was diminished to the level that is typically evoked by a monocular stimulus while the superficial layers exhibited significant inhibition. This order of excitement and inhibition within V1’s laminar microcircuit suggests that cortico-cortical interactions mediate interocular suppression within V1, with each portion of the microcircuit playing a distinct role. These findings provide important new information for neuronal models of binocular processing (Bridge and Cumming, 2008; Bhaumik and Shah, 2014).

## Author Contributions

M.A.C., K.D., and A.M. conceptualized and designed the study. M.A.C., K.D., and M.S. trained animals and implemented the experiments. A.M., M.A.C., J.A.W., and K.D. collected and analyzed neurophysiological data. M.A.C. prepared figures for publication. A.M. and M.A.C. wrote the manuscript with input from other authors.

## Acknowledgments

The authors would like to thank S. Amemori, Dr. T. Apple, M. Feurtado, K. George-Durrett, A. Graybiel, N. Halper, P. Henry, M. Johnson, Dr. C. Jones, M. Maddox, L. McIntosh, Dr. A. Newton, J. Parker, C. Thompson, K. Torab, C. Subraveti, B. Williams, and R. Williams for technical advice and assistance; R. Krauzlis, F. Tong, and K. Dieter for comments on previous drafts of this manuscript. This work was supported by a research grant from the National Eye Institute (1R01EY027402-01). K.D. and J.A.W. are each supported by a National Eye Institute Training Grant (5T32EY007135-23). A.M. is supported by a research grant of the Whitehall Foundation, a Career Starter grant by the Knights Templar Eye Foundation, and a Fellowship of the Alfred P. Sloan Foundation. The authors declare no competing financial interests.

